# High-fidelity disentangled cellular embeddings for large-scale heterogeneous spatial omics via DECIPHER

**DOI:** 10.1101/2024.11.29.626126

**Authors:** Chen-Rui Xia, Zhi-Jie Cao, Ge Gao

**Author notes:** Authors to whom correspondence should be addressed (for G.G.), or (for Z.J.C.). These authors contributed equally to this work (Chen-Rui Xia, Zhi-Jie Cao).

## Abstract

The functional role of a cell, shaped by the sophisticated interplay between its molecular identity and spatial context, is often obscured in current spatial modeling. Aiming to model large-scale heterogeneous spatial data *in silico* properly, DECIPHER produces high-fidelity disentangled embeddings, not only achieving superior performance in systematic benchmarks, but also empowering various real-world applications. We further demonstrated that DECIPHER is scalable to atlas-scale datasets, enabling global analysis which is largely infeasible to current state-of-the-arts.

## Main

In multicellular organisms, cells must interact to organize into three-dimensional tissue structures. The spatial context of an individual cell is as critical as its intrinsic properties in determining its physiological function and potential pathological changes^1,2^. While spatial omics technologies enable systematic characterization of both of these features, their proper *in silico* representation remains a serious challenge^3–7^.

Embedding, or the procedure of projecting original high-dimensional spatial data as low-dimensional vectors, has been widely adopted to enrich biologically relevant signals while simultaneously correcting or removing noise and technical artifacts^8,9^. Inspired by its great success in single-cell omics^7^, the current practice in spatial omics is to generate a holistic embedding with information of both the cellular transcriptome and spatial location encoded into one single low-dimensional space^3–5^. While this approach eases model design and implementation, it effectively introduces intrinsic entanglement between the intra-cellular molecular identity and inter-cellular context, obfuscating the dissection of their interplay which is rather important for many biological applications like cell-cell communication and signaling-induced cell localization (Fig. 1a).

**Fig. 1:**
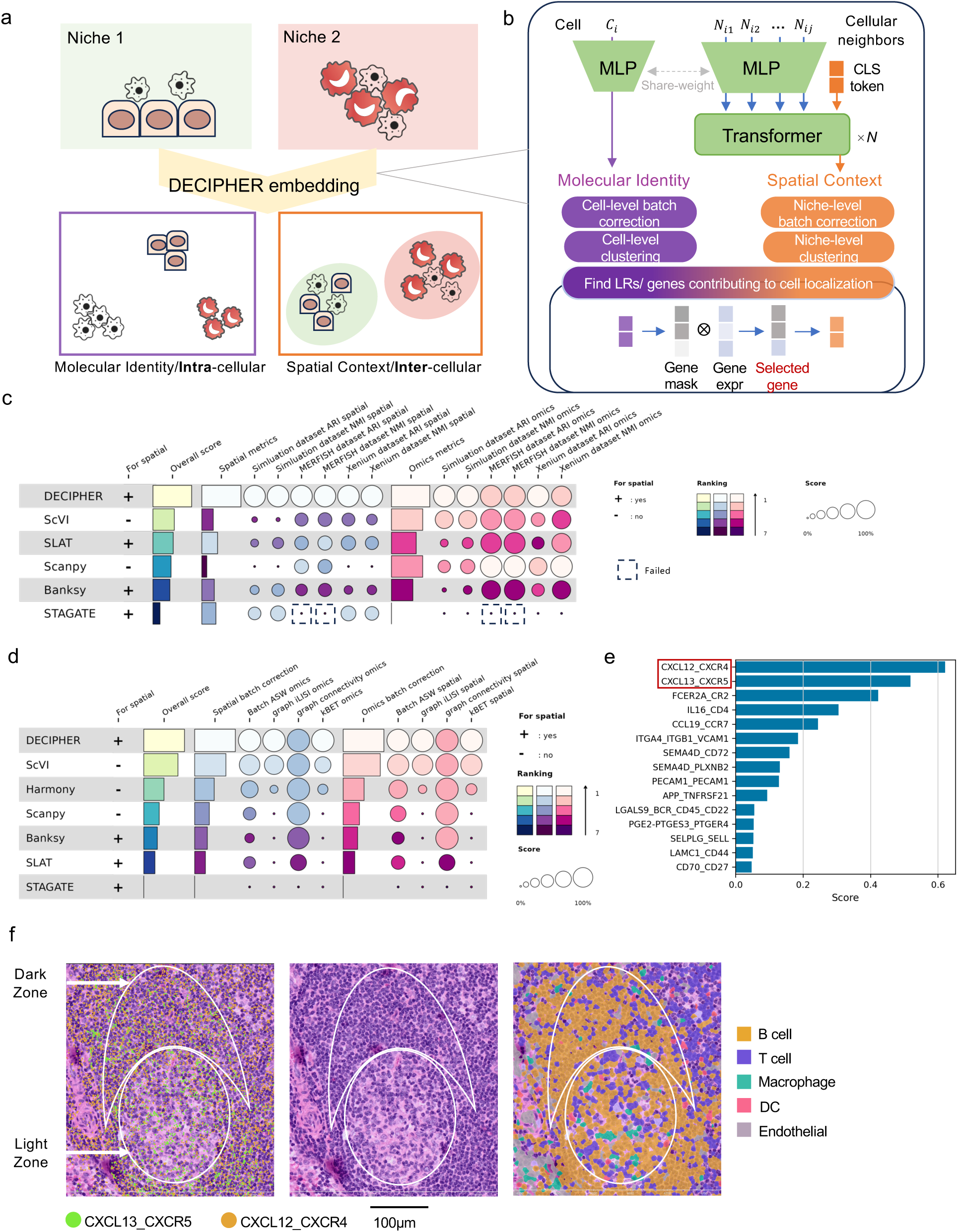
DECIPHER for learning disentangled high-fidelity intra- and inter-cellular embeddings. **a,** DECIPHER models the intracellular molecular identity and intercellular spatial context of cells, respectively. Cells with similar omics profiles are assigned similar molecular identity embeddings, while cells located inside the same niche receive similar spatial context embeddings. **b,** Architecture of DECIPHER, which uses an omics encoder (MLP) to obtain the molecular identity embedding of cell *C_i_* and uses a spatial encoder (transformer) to obtain the spatial context embedding of the cell from *C_i_*’s s spatial neighbors (*N_i1_* … *N_ij_*). The bottom buttons show applications based on identity or/and context embeddings. In particular, to find LR pairs/genes contributing to cell localization, a LR/gene level mask is learned from the molecular identity embedding by reconstructing the corresponding spatial context embedding via the selected gene expression (see Extended Data Fig. 3a and 3b for details). **c,** Summary of benchmark results. Normalized mutual information (NMI) and adjusted rand index (ARI) were reported against the original spatial region annotations (termed “Spatial metrics”) and against cell type annotations (termed “Omics metrics”) in three spatial datasets (Simulation, MERFISH and Xenium) (**Methods**). The “overall score” is the average of all metrics. STAGATE failed to run on the MERFISH dataset because of GPU memory overflow (capping at 80 GB); details are reported in Extended Data Fig. 3a and 3c. **d,** Summary of batch correction benchmark results. Batch ASW (average silhouette width), batch GC (graph connectivity), batch iLSI (integration local inverse Simpson’s index), and batch kBET (*k*-nearest-neighbor batch effect test) were calculated against spatial pattern annotations (termed “Spatial batch correction”) and cell type annotations (termed “Omics batch correction”) (**Methods**). The “overall score” is the average of all metrics. Details are reported in Extended Data Fig. 3b. **e,** Bar plot showing the top 10 most important LR pairs accounting for B-cell localization in the lymph node dataset calculated by the LR selection model in Extended Fig. 3a. **f,** Visualization of a typical GC; the dark zone and light zone were annotated by an anatomical expert. The left panel shows the CXCL13_CXCR5 (green) and CXCL12_CXCR4 (yellow) signal strengths (**Methods**). The middle panel shows the original H&E-stained image of the GC. The right panel shows the cell type, with colors representing different cell types.

Here, we present DECIPHER, a context-aware deep learning model for effectively and efficiently disentangling cellular embeddings for large heterogeneous spatial slices with high fidelity (Fig. 1a). Briefly, DECIPHER consists of two interconnected components: the “Omics Encoder”, for learning an intracellular molecular identity-oriented embedding from the expression profile, and the “Spatial Encoder”, for projecting the cell’s neighborhood context into an independent spatial context embedding space (Fig. 1b, **Methods**). Both components were optimized simultaneously via a dedicated cross-scale contrastive learning procedure, directly modeling cellular disentangled representations via self-attention-based transformation (Extended Data Fig. 1, **Methods**).

To evaluate the quality of the resulting embeddings, we first compared them with the results of several state-of-the-art methods applied to two well-curated real-world datasets (the Xenium breast cancer dataset^10^ and the MERFISH brain dataset^11^) and a synthetic dataset with clear ground truths for cell types, spatial domains and batch effects (**Methods**, Supplementary Table 1). Across all three datasets, DECIPHER’s spatial context embedding achieved superior performance for both spatial domain identification^3–5^ (Fig. 1c Extended Data Fig. 2a) and batch correction (Fig. 1d and Extended Data Fig. 2b). Notably, while DECIPHER has comparable performance to that of state-of-the-art scRNA-seq embedding methods in the MERFISH and Xenium datasets, where the gene panels include only a few hundred genes, it is superior in the synthetic dataset covering the whole transcriptome (**Methods**, Fig. 1c and Extended Data Fig. 2c), suggesting a higher performance potential that could be realized through further improvements in spatial omics data quality. Notably, existing spatial omics embedding models generally exhibit subpar performance in terms of cell type identification, likely due to resolution degradation from the holistic embedding strategy, which DECIPHER avoids by design (Fig. 1c, Extended Data Fig. 2a and 2c).

The high-fidelity disentangled embeddings provided by DECIPHER enable direct modeling of sophisticated interactions between intra-and extra-cellular factors, which is otherwise impossible with a holistic modeling approach. For instance, by reconstructing the spatial context embedding with a ligand-receptor (LR)-level binary mask vector learned from each cell’s intracellular embedding, we could identify key cell-cell communication (CCC) molecules contributing to cellular localization (Fig. 1b, Extended Data Fig. 3a and 3b, **Methods**). For example, DECIPHER successfully identified CXCL12_CXCR4 and CXCL13_CXCR5 as the two most important LR pairs accounting for the localization of maturing B cells within germinal centers (GCs)^12^ (Fig. 1f, Extended Data Fig. 3c-3f, **Methods**). Previous studies have demonstrated that CXCR4 is crucial for guiding B cells to the dark zone and that CXCR5 is crucial for guiding B cells to the light zone^13,14^. Further visualization via paired H&E images revealed enrichment of CXCR13_CXCR5 signaling in the light zone and CXCR12_CXCR4 in the dark zone (Fig. 1f), further indicating that DECIPHER correctly identified critical LR pairs determining B-cell localization in GCs. Notably, canonical holistic embedding-based methods would miss the CXCL13_CXCR5 signal due to their lower activity levels (Extended Data Fig. 4a-d), highlighting the high resolution of DECIPHER’s disentangled embeddings.

Of note, the DECIPHER’s high-fidelity embeddings further empower calling key genes over technologies with restricted gene coverage (Fig. 1b and Extended Data Fig. 3a, **Methods**). As demonstrated in a Xenium breast cancer dataset, lipocalin-type prostaglandin D2 synthase (*PTGDS*) emerged as the most relevant gene for the localization of both T and B cells (Fig. 2a). T and B cells expressing high levels of *PTGDS* were either enriched at the tumor interface or found infiltrating the tumor (Fig. 2b). A previous breast cancer study using IMC imaging independently identified *PTGDS* as a crucial marker for infiltrating lymphocytes (CD4+/CD8+/CD19+)^15^, confirming our findings. Among the remaining top-ranked genes, *CXCL12*^16^, *CD86*^17^, *SFPR4*^18^ and *CXCR4*^19^ have also been linked to lymphocyte activation and recruitment (Extended Data Fig. 5a and 5b). On the contrary, the same differential expression analysis based on canonical holistic embedding just failed to identify *PTGDS* and others (Extended Data Fig. 5c-e, 6f, and **Methods**).

**Fig. 2:**
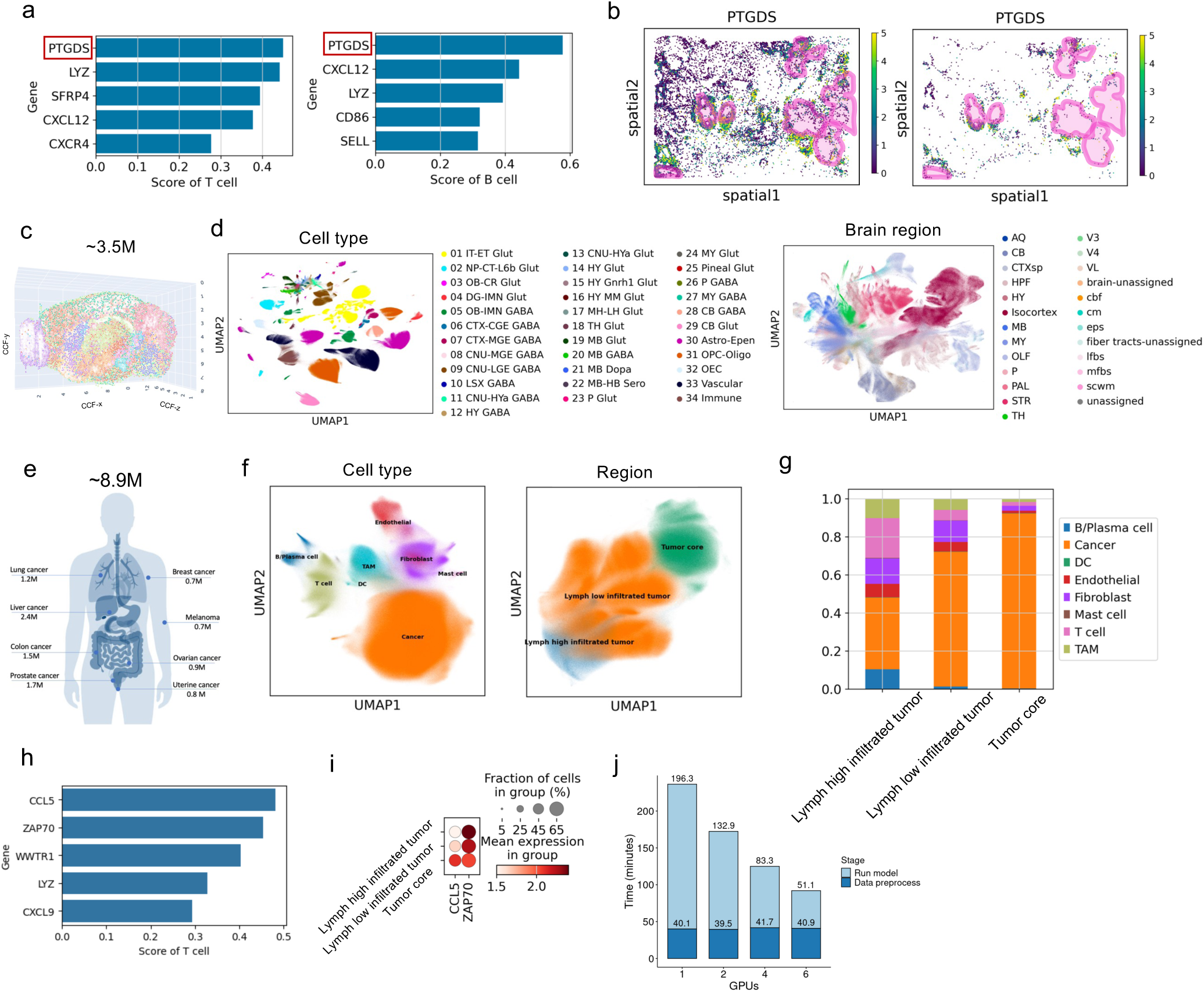
Applications based on DECIPHER embedding. **a,** Bar plot showing the top 5 genes of T cells (left panel) and B cells (right panel) identified by the gene selection model (described in Fig. 1e) in the breast cancer dataset. **b,** Spatial expression of *PTGDS* in T cells (left panel) and B cells (right panel). Pink circles indicate the DICS tumor region annotated by the original author. **c,** Diagram illustrating the mouse brain 3D atlas in CCF coordinates^38^. **d**, UMAP visualization of the molecular identity embeddings (left panel) and spatial context embeddings (right panel) of DECIPHER on the 3D mouse brain atlas. **e,** Diagram illustrating the cancer types of the human pan-cancer spatial atlas. **f,** UMAP visualization of the intracellular identity embeddings (left panel) and spatial context embeddings (right panel) of DECIPHER on the human pan-cancer spatial atlas. **g,** Cell type composition in different spatial regions defined in **f**. **h,** Bar plot showing the top 5 genes of T cells identified by the gene selection model in the human pan-cancer atlas. **i,** Expression of *CCL5* and *ZAP70* in T cells across spatial regions defined in **g**. **j,** Time consumed with different numbers of GPUs. Astro, astrocyte; AQ, aqueduct of Sylvius, CB, cerebellum; CGE, caudal ganglionic eminence; CNU, cerebral nuclei; CR, Cajal–Retzius; CT, corticothalamic; CTX, cerebral cortex; CTXsp, cortical subplate; DG, dentate gyrus; EA, extended amygdala; Epen, ependymal; EPI, epithalamus; ET, extratelencephalic; GC, granule cell; HB, hindbrain; HPF, hippocampal formation; HY, hypothalamus; HYa, anterior hypothalamic; IMN, immature neurons; IT, intratelencephalic; L6b, layer 6b; LGE, lateral ganglionic eminence; LH, lateral habenula; LSX, lateral septal complex; MB, midbrain; MGE, medial ganglionic eminence; MH, medial habenula; MM, medial mammillary nucleus; MY, medulla; NP, near-projecting; OB, olfactory bulb; OEC, olfactory ensheathing cells; OLF, olfactory areas; Oligo, oligodendrocytes; OPC, oligodendrocyte precursor cells; P, pons; PAL, pallidum; STR, striatum; TH, thalamus; V3, third ventricle; V4, fourth ventricle; VL, ventral lateral nucleus; CM, centromedian nucleus; EPS, entorhinal-perirhinal-solitary complex; IFBS, internal fiber bundles; MFBS, medial forebrain bundles; SCWM, sub-cortical white matter. Neurotransmitter types: DOPA, dopaminergic; GABA, GABAergic; Glut, glutamatergic; Nora, noradrenergic; Sero, serotonergic^39^.

Technological advances have enabled the generation of spatial omics datasets exceeding millions of cells^20^. For example, a mouse brain atlas comprising 151 consecutive slices of a single brain with 3.5 million cells was constructed to capture the 3D organization of the central nervous system^21^ (Fig. 2c, Extended Data Fig. 6a); this dataset exceeds the scalability of existing computational methods, even if allocated 1 TB system memory, 80 GB GPU memory and 48 hours of runtime. A major design consideration of DECIPHER is its robustness and scalability, allowing its application to the entire brain atlas. Its molecular identity embeddings match well with the original cell type annotations (Fig. 2d, left panel), whereas the spatial context embeddings correspond accurately to brain anatomical regions (Fig. 2d, right panel, Extended Data Fig. 6b). Interestingly, when only 2D coordinates of cells within slices were used and 3D information was omitted, the quality of the spatial context embeddings was significantly reduced, and even basic brain regions were not distinguished (Extended Data Fig. 6c), underscoring the importance of 3D information for uncovering complex tissue spatial patterns^22^.

DECIPHER’s advantages in terms of both performance and scalability enable global analysis for larger spatial atlases. The human pan-cancer spatial atlas consists of 8.9 million cells spanning eight different cancer types^23^ (Fig. 2e), with significant batch effects across slices from various cancer types (Extended Data Fig. 7a). DECIPHER successfully distinguished cell types in the molecular identity embeddings, producing high-quality batch-corrected spatial context embeddings that facilitate the comparison of spatial patterns across cancer types (Fig. 2f, Extended Data Fig. 7b). Notably, spatial patterns in cancer typically exhibit continuous variation in cell type density, reflecting the dynamic nature of tumor growth, invasion, and lymphocyte infiltration (Extended Data Fig. 7c). Thus, the tumor microenvironment (TME) can be roughly classified into three regions on the basis of the density of different cell types^24^ (Fig. 2f and 2g): the tumor core (dominated by tumor cells), lymph-low infiltrated tumor, and lymph-high infiltrated tumor. We found that T and B cells were significantly colocalized in the lymph-high infiltrated tumor regions (*p* value < 10^-16^, Extended Data Fig. 7c and 7d), which was consistent with their formation of tertiary lymphoid structures (TLSs) in microscopy images (Extended Data Fig. 7e). We further identified the genes most associated with T-cell localization using the same approach as described above (**Methods**) and identified *CCL5* and *ZAP70* as the top two genes (Fig. 2h). Interestingly, we observed that T cells with high *CCL5* expression tend to reside in the tumor core, whereas T cells with high *ZAP70* expression are more likely to be excluded (Fig. 2i). *ZAP70* is a key factor in T-cell activation and response to antibodies^25^ but may be suppressed within tumors^26^. *CCL5* is known to induce Treg infiltration in tumors and restrain T-cell activation^27^. These factors may contribute to the highly exhausted state observed in tumor core-residing T cells, which exhibit high expression of immunosuppressive checkpoint genes, including *PD-1*, *LAG-3*, and *CTLA-4*. This aligns with the “terminally exhausted T cells” (Tex terminal, characterized by PD-1+, LAG3+, CTLA4+, GZMA+, GLNY+, GZMB+ and IFNG+), as defined in a nonspatial pan-cancer T-cell study^28^ (Extended Data Fig. 7f). Our findings highlight a strong association between the T-cell state and spatial localization in the global TME, which is difficult to observe in single spatial slices because of the limited fields of view (Extended Data Fig. 8). In contrast, other state-of-the-art methods fail to process entire datasets and produce suboptimal results on subsampled data (Extended Data Fig. 9).

Aiming to model spatial omics data *in silico* properly, DECIPHER has demonstrated superior performance in various benchmarks and real-world case studies. We anticipate that with continued technological advances, high-quality spatial atlases covering entire organs or even organisms will eventually emerge. In addition to its unique ability to produce high-fidelity embeddings, DECIPHER’s contrastive self-supervised learning strategy combined with the transformer architecture is well suited for large-scale heterogeneous spatial pretraining (Fig. 2j). Once such organism-scale data become available, DECIPHER could serve as the backbone of a pretrained spatial foundation model, providing a reliable starting point for addressing various biological questions involving cellular functions within native contexts.

## Methods

### Framework

DECIPHER requires one or multiple single cell spatial omics datasets consisting of the raw expression matrix and spatial coordinates of cells (supporting 2D or 3D coordinates). At the omics level, the raw expression matrix undergoes standard processing^29^, including log-based normalization, highly variable gene selection, and scaling. At the spatial level, for each cell *i*, *k*-nearest neighbors in the spatial coordinate space are considered the spatial neighbors of cell *i*, denoted as *𝒩_i_* (with *k* defaulting to 20).

The DECIPHER model is composed of an “omics encoder” and a “spatial encoder” (Fig. 1b), which are used to learn molecular identity and spatial context embeddings, respectively. Specifically, the “omics encoder” is a multilayer perceptron (MLP) that projects high-dimensional single-cell expression data into the molecular identity latent space (by default, 32 dimensions), denoted as 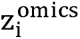. The “spatial encoder” adopts a transformer architecture^30,31^ (by default, 3 layers with 1 attention head). It takes the omics features of all spatial neighbors of *cell_i_* (excluding *cell_i_* itself) and a trainable token (CLS token) as input tokens. The output embedding of the CLS token is ultimately used as the spatial context embedding 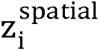 of *cell_i_* (by default, 32 dimensions). To reduce computational complexity, the input tokens are not high-dimensional expressions but rather latent representations learned by the omics encoder.

### Training process

The DECIPHER model is optimized through novel multiscale contrastive learning, for which the construction of positive sample pairs is crucial. In contrast to the data augmentation strategies used in the field of computer vision, such as random cropping and color inversion^32^, we simulate dropout events to obtain augmented views of single-cell omics data. Specifically, for each cell, we randomly mask genes twice on the basis of probability *p* (defaulting to 0.5) to obtain two different “dropout views” of the same cell. Contrastive loss is used to move the two dropout views of the same cell close to each other, while views of different cells are pushed apart^32^. Data augmentation occurs only during training. The complete omics feature of each cell is used directly as the input in the inference stage.

Additionally, to handle batch effects across multiple slices, we include mutual nearest neighbor (MNN) pairs across slices as extra positive pairs in addition to the positive pairs obtained through the above data augmentation strategy.

DECIPHER utilizes the NT-Xent contrastive loss function^32^:

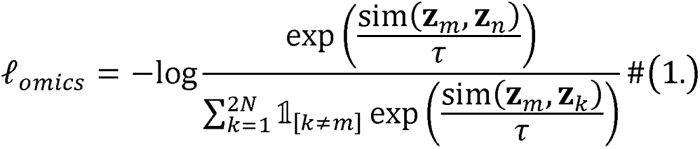

Here, *z_m_* and *z_n_* are two views of the same cell, *z_m_* and *z_n_* are two views of different cells, *N* is the batch size, and τ is the temperature coefficient.

For spatial context embedding, since each neighbor of cell *i* has two “dropout views”, we naturally obtain two views of the spatial context embedding of cell *i* via the spatial encoder. We also use the NT-Xent loss to encourage positive spatial context embedding pairs to be close to each other and to push negative pairs apart (Fig. 1b).

Finally, the total loss of the model is calculated as the average of the contrastive losses at the omics and spatial compartments:

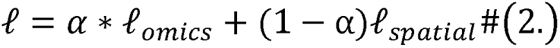

where α is 0.5 by default. DECIPHER is built on the PyTorch framework and supports multi-GPU parallelization. By default, DECIPHER first warms up the omics encoder for 3 epochs with a learning rate of 0.01 and then trains the omics encoder and spatial encoder synergistically for 5 epochs or 10,000 steps with a decayed learning rate of 0.00001.

We tested DECIPHER’s robustness to key hyperparameters including (1) the latent space dimension, (2) the layer of the transformer, (3) the number of attention heads, (4) the neighbor size *k*, and (5) the loss weight α; the results are reported in Supplementary Table 2.

### Localization-related gene/ligand–receptor pair selection model

For the gene selection model, we used a parameterized neural network *t* to quantify the importance of each gene for the spatial context embedding of cell *i* and then binarized it into a differentiable mask via Gumbel-sigmoid sampling^33^. The Hadamard product is performed with the log-normalized original expression levels of the cells to obtain the selected genes only:

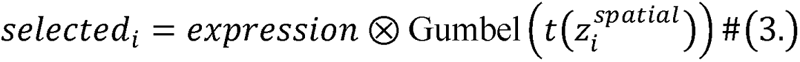

*selected_i_* contains only the selected gene set 𝒮 of cell *i* (Fig. 1e).

The training objective is to identify the gene set 𝒮 most related to the spatial context embedding. We transformed this problem into reconstructing the spatial context embedding similarity graph. Specifically, we first construct a *k*-nearest neighbors (*k*-NN) graph *𝒢_Spatial_* on the basis of spatial context embedding (using “scanpy.pp.neighbors()”). During training, we use *selected_i_* to reconstruct the edges of 𝒢*_Spatial_* (Extended Data Fig. 3a), with the loss function being the negative log-likelihood of edge reconstruction:

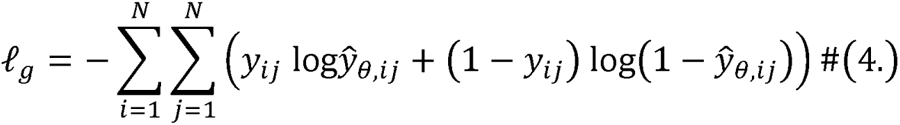

In the above equation, *Y_ij_* indicates whether there is an edge between *cell_i_* and *cell_i_* in *𝒢_Spatial_*. Additionally, to filter out irrelevant genes, we add an extra L_1_ regularization to the size of 𝒮. The overall loss is written as:

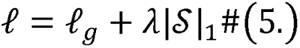

λ is set to 1 by default. During training, the model applies back propagation to learn the gene mask, gradually filtering out genes that do not contribute to predicting the spatial context embedding. By default, the model is trained for 300 epochs with a learning rate of 0.01. After training is complete, the resulting 𝒮*_i_* represents the key genes for the spatial positioning of cell *i*. Then, for each cell type, we determine the importance of gene *g* for the cell type by dividing the number of times each gene is selected (*X_g_*) by the size of the cell type (*N*):

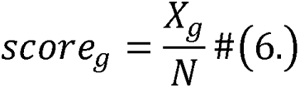

For the ligand-receptor selection model, the basic units are not genes but rather ligand-receptor pairs. Additionally, we incorporate an LR mask that focuses exclusively on the following:

1. The receptor genes expressed by the cell; and
2. The ligand-receptor pairs where at least one neighboring cell within a 100 µm radius expresses the ligand gene (100 µm represents a typical cell-cell interaction distance^34^).

Other settings are consistent with the gene selection model above. Note that in this model, we do not consider the cell type of the sender (ligand) because, for the receptor cell, the sender of the signal is not visible. The activity of each ligand-receptor pair in cell *i* is defined as the product of expression between the receptor genes *e^receptor^* in cell *i* and the sum of the ligand gene expression *e^ligand35r^* within the 100 µm radius (denoted as *𝒩_n_*):

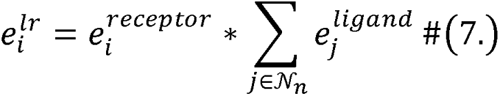

### Benchmark

#### Public datasets

The MERFISH brain dataset^11^ and Xenium breast cancer dataset^10^ are publicly available. The MERFISH brain dataset contains 378,918 cells and 374 genes, and the Xenium breast tumor dataset contains 164,079 cells and 313 genes. We use the cell type and spatial domain annotations from the original publications.

#### Simulation dataset

We simulated a spatial dataset in which the ground truths of the cell types and spatial patterns were clear. Specifically, we selected one 10x 5’ and one 10x 3’ based PBMC dataset from the official 10x datasets from the same donor. These two datasets contain technical batch effects (Extended Data Fig. 10a). For each dataset, we annotated cell types via the same criteria, retaining only three distinct cell types: T cells (CD3D), B cells (CD79A), and monocytes (CD14, CD16) (Extended Data Fig. 10b).

The two datasets contained 4,512 cells and 3,893 cells (8,405 in total), respectively, and 31,915 shared genes.

Then, we simulated three identical spatial patterns for both datasets (Extended Data Fig. 10c):

1. Pattern 1: Monocytes and T cells randomly mixed.
2. Pattern 2: Only T cells, randomly distributed.
3. Pattern 3: B cells and T cells randomly mixed.

Thus, the two datasets are homogeneous in terms of cell types and spatial patterns.

#### Methods

The benchmark methods Scanpy, Harmony, scVI, Banksy, STAGATE, and SLAT were executed via the Python packages “Scanpy” (v1.10.1), “harmonypy” (v0.0.6), “scvi-tools” (v1.1.2), “Banksy_py” (last commit 10770c2), “STAGATE_pyG” (latest commit 8b9c8ef), and “scSLAT” (v0.2.1), respectively, in Python (v3.11). For each method, we used the default data preprocessing steps recommended by the original authors.

#### Metrics

For identity and context embedding, we used two common metrics: the adjusted rand index (ARI) and normalized mutual information (NMI). For batch effect removal, we used four common metrics: batch ASW, batch GC, batch iLSI, and batch kBET. All metrics were calculated following the standard procedures in scIB^36^. All benchmark tasks were allocated 8 CPU cores (Intel Xeon Platinum 8358), 128 GB of memory, and one GPU (A100-80G) by Slurm.

### Case studies

#### Mouse brain 3D atlas profiling

The mouse brain atlas consists of ∼3.5 M quality-controlled cells with 1,122 detected genes, derived from 151 consecutive MERFISH slices of a single adult mouse brain^21^, including detailed annotations of cell types and spatial domains. These slices are also registered to 3D CCF coordinates. We trained DECIPHER using 3D CCF coordinates or the original 2D slice coordinates (Fig. 3b and 3d). We calculated the density of each cell type on UMAP via the “scanpy.tl.density()” function (Fig. 3c).

For the other methods, we randomly down sampled the dataset to 200,000 cells (the maximum size for STAGATE) and subsequently ran each model with its default parameters.

#### Human pan-cancer spatial atlas profiling

The human pan-cancer spatial atlas includes 16 slices from 8 cancer types (2 colon cancer, 2 liver cancer, 2 melanoma, 4 ovarian cancer, 2 prostate cancer, 2 lung cancer, 1 breast cancer, and 1 uterine cancer), containing a total of 8,696,580 cells and 500 genes, analyzed using the MERSCOPE platform. We manually annotated the major cell types in the pan-cancer MERSCOPE data via the following classical markers: T cells (*CD3D*), cancer cells (*VCAM1*, *MKI67*), and B cells (*CD79A*).

Owing to the significant batch effects between slices, we used the MNN batch effect removal option in DECIPHER (as mentioned above). The runtime included both data processing and algorithm execution, which were performed on a server equipped with 8 A100-80G GPUs. We calculated the density of each cell type in UMAP via the “scanpy.tl.density()” function (Fig. 3c).

For the other methods, we randomly down-sampled the dataset to 200,000 cells (the maximum size for STAGATE in our server) and subsequently ran each model with its default parameters.

#### Identification of key localization-related genes in the Xenium breast tumor dataset

The Xenium breast tumor dataset contains 164,079 cells and 313 genes and was annotated by the original authors. We ran DECIPHER on this dataset with its default parameters. On the basis of DECIPHER embedding, the gene selection model was run separately for T cells and B cells. For each cell type, genes were ranked according to their importance scores; the top 5 genes are presented.

For comparison, we clustered the spatial context embeddings from DECIPHER, Banksy, and SLAT to obtain spatial domains and then identified DEGs between tumor domains and nontumor domains using the “scanpy.tl.rank_genes_groups()” function.

#### Identification of key CCC molecules contributing to cellular localization in the Xenium 5k data

The Xenium 5k lymph node dataset includes 708,983 cells and 4,624 elaborately selected genes^37^, covering important cell-cell communication and signaling pathways. The corresponding cell segmentation and H&E-stained images were provided by 10x Genomics. First, we annotated the major cell types in the Xenium 5k dataset according to the following classical markers (Extended Data Fig. 3c and 3d): T cells (*CD3E*), dendritic cells (DCs, *ITGAX*), B cells (*CD79A*), endothelial cells (*PLVAP*), macrophages (*MMP9*), plasma cells (*MZB1*), follicular dendritic cells (fDCs, *CR2*), plasmacytoid dendritic cells (pDCs, *IRF7*), and vascular smooth muscle cells (VSMCs, *NOTCH3*). A group of cells with significantly lower counts were annotated as “low_quality”. The activity of each LR pair is defined following Formula (7).

DECIPHER was run with the default parameters. Molecular identity embedding from DECIPHER corresponded well with the cell types, and spatial context embedding produced 10 Leiden clusters with 0.5 resolution (Extended Data Fig. 3f).

We ran the ligand-receptor selection model for B cells. Ligand-receptor pair data were sourced from the latest version of CellChatDB^35^ (http://www.cellchat.org/cellchatdb/), which contains 3,124 LR pairs, among which 1,078 pairs were detected in this dataset. We used the product of ligand and receptor gene expression. For visualization, a pathologist selected a typical GC field and annotated the light zone and dark zone in the GC. We visualized the strength of the CXCR5_CXCL13 and CXCR4_CXCL12 signaling pathways and the cell type information within the GC.

For comparison, we clustered the embeddings from Banksy and SLAT to obtain spatial domains and then identified differentially expressed LRs between the GC and non-GC regions using the “scanpy.pp.rank_genes_groups()” function.

## Data available

All datasets used in this study have already been published and were obtained from public data repositories. Supplementary Table 1 provides the associated publications and download URLs.

## Code available

DECIPHER is available at https://github.com/gao-lab/DECIPHER, along with code and detailed tutorials for reproducibility.

## Acknowledgments

We thank Mr. Z. Qi for his assistance with H&E image annotation, as well as Drs. Z. Zhang, L. Tao, F. Tang, X.S. Xie, C. Li, and J. Lu at Peking University for their helpful discussions and comments during the study. This work was supported by funds from the National Key Research and Development Program of China (2022ZD0115004), as well as the State Key Laboratory of Protein and Plant Gene Research, the Beijing Advanced Innovation Center for Genomics (ICG) at Peking University, the Changping Laboratory, and the Shaw Foundation Hong Kong Limited. The research of C.R.X. is supported by the National Natural Science Foundation of China (grant no. 323B2017). The research of Z.J.C. is supported by the China Postdoctoral Science Foundation (grant no. 2023T160009 to Z.J.C.). The research of G.G. was supported in part by the National Program for Support of Top-Notch Young Professionals.

## Author contributions

G.G. and Z.J.C. conceived the study and supervised the research. C.R.X. and Z.J.C. designed and implemented the computational framework and conducted benchmarks and case studies with guidance from G.G. C.R.X., Z.J.C., and G.G. wrote the manuscript.

## Figure legends

**Extended Data Fig. 1:**
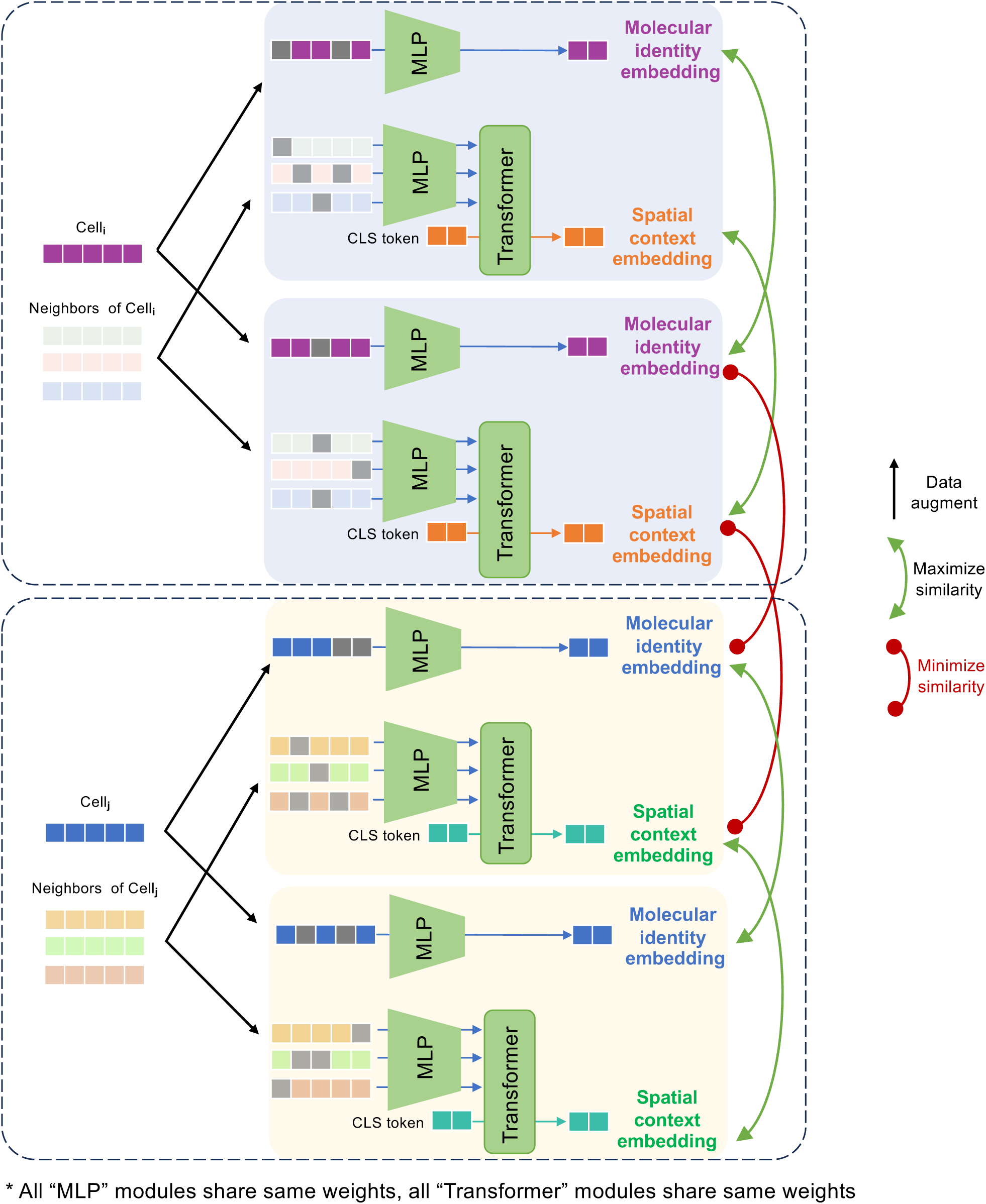
DECIPHER optimization process. Detailed optimization process of DECIPHER. Two augmented views are obtained for each cell through random dropout (**Methods**). The two views are processed separately by the DECIPHER model, and two molecular identity embedding views and two spatial context embedding views are generated. During training, the model maximizes the similarity between the two views of the same cell while minimizing the similarity between the views of different cells.

**Extended Data Fig. 2:**
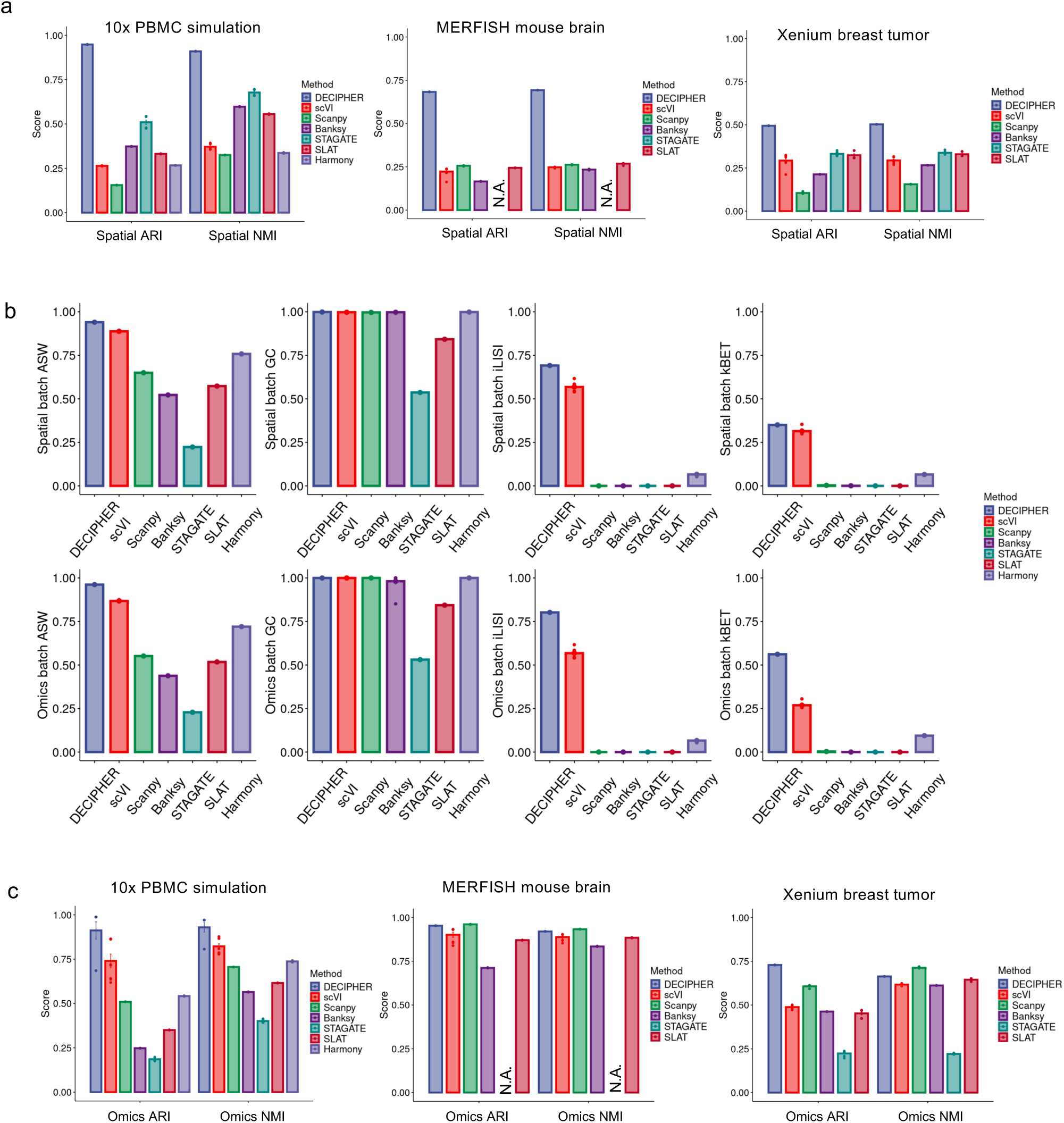
Benchmark results. **a,** Benchmark results for spatial clustering. NMI and ARI were reported against the original spatial region annotations. From left to right are results on the 10x mimic dataset, the MERFISH brain dataset and the Xenium breast tumor dataset. **b,** Benchmark results for batch correction. batch ASW, batch GC, batch iLSI, and batch kBET were calculated against spatial region annotations (first row) and cell type annotations (second row), respectively. **c,** Benchmark results for single-cell clustering. NMI and ARI were reported against the original cell type annotations. From left to right are the results on the 10x mimic dataset, the MERFISH brain dataset and the Xenium breast tumor dataset. STAGATE failed to run on the MERFISH brain dataset because of GPU memory overflow (capping at 80 GB). *n* = 8 repeats with different random seeds. The error bars indicate the means ± s.d.

**Extended Data Fig. 3:**
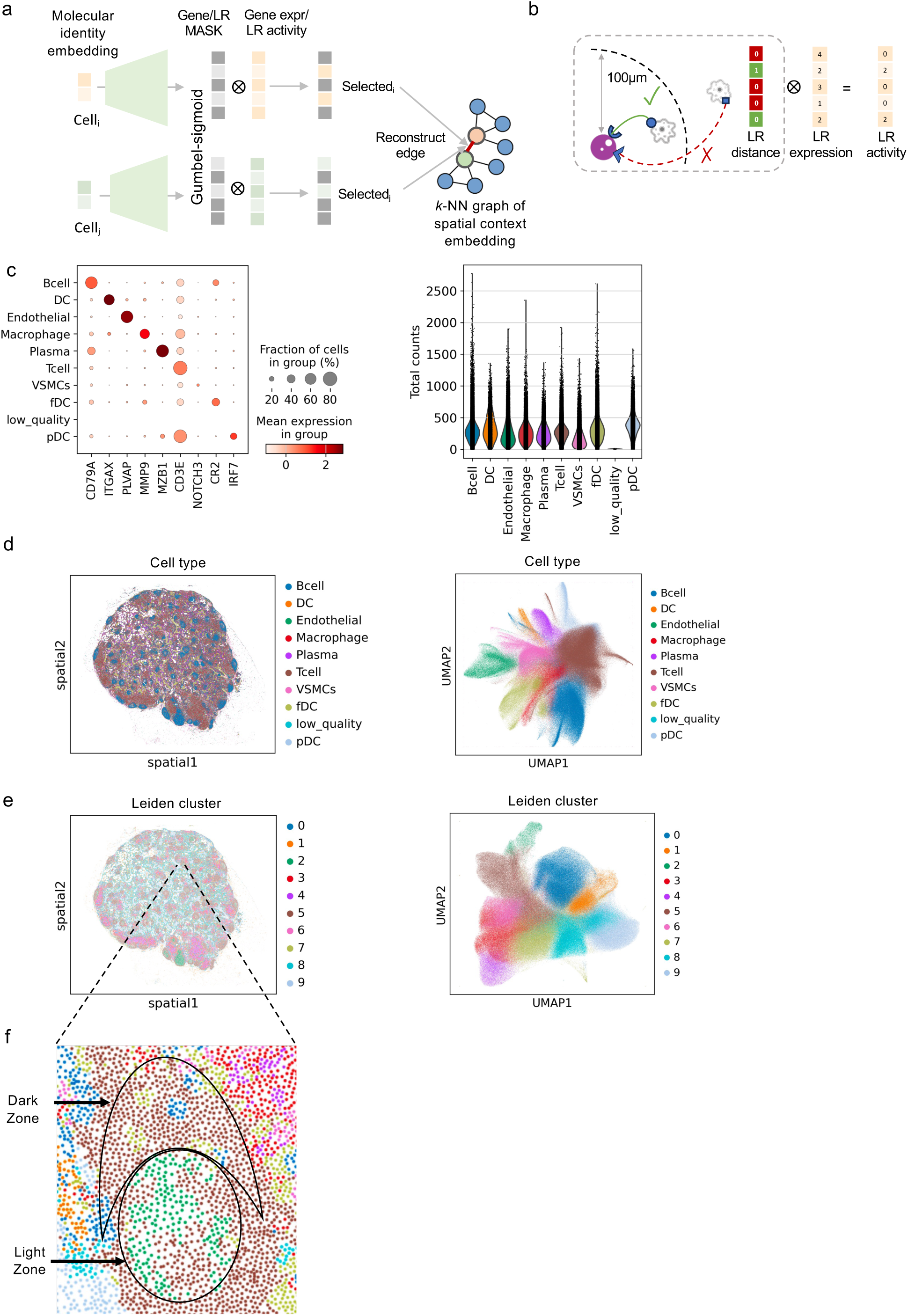
Profiling the lymph node dataset via DECIPHER. **a,** Detailed illustration of the LR/gene selection model. This model uses a neural network and a Gumbel-sigmoid layer to generate learnable LR/gene selection masks from the molecular identity embedding. The target of training the model is to reconstruct the spatial embedding k-NN graph (**Methods**). **b,** Illustration for LR activity calculation, which is the production of LR expression^35^ (**Methods**) and an extra LR mask: if cell *i* expresses the receptor gene of one LR pair and cell *i*’s neighbors within 100 µm express the corresponding ligand gene, the corresponding value in the mask is set to “1” (green); otherwise, it is set to “0” (red). **c,** Expression of marker genes in each cell type (left panel); a group of cells were annotated as ‘low_quality’ due to the low detected counts (right panel). **e,** Visualization of cell types on spatial slices and UMAP plot of DECIPHER’s molecular identity embedding. **d,** Visualization of the Leiden cluster defined by DECIPHER’s spatial context embedding on a spatial slice (left panel) and the corresponding UMAP (right panel). **e,** Magnification of the germinal center region shown in Fig. 2g on spatial slices, colored by the Leiden cluster in **e**.

**Extended Data Fig. 4:**
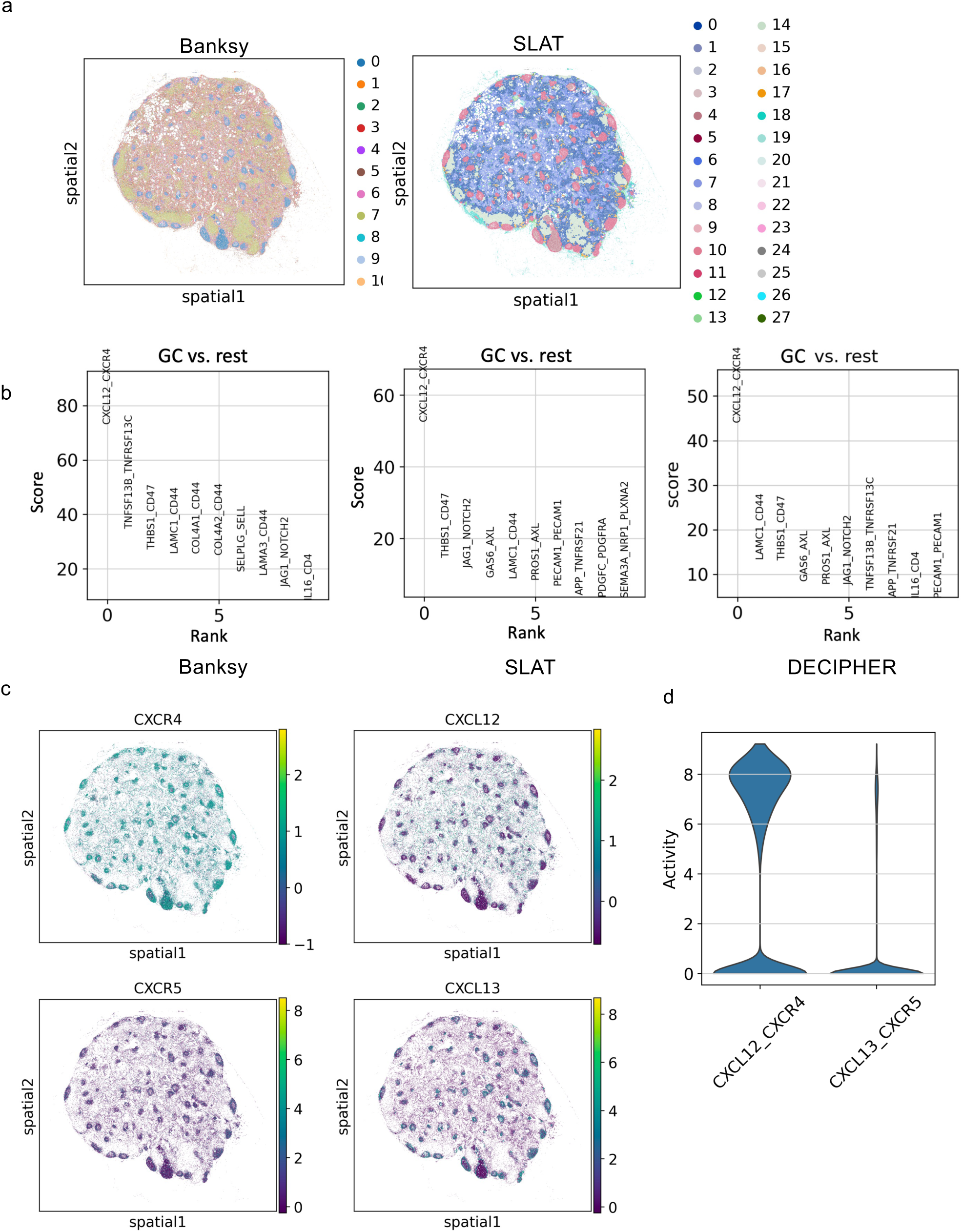
Comparison with baseline methods in the lymph node dataset. **a,** UMAP visualization and spatial visualization based on Banksy and SLAT embedding, colored according to the Leiden clustering results. STAGATE failed because of GPU memory overflow (capping at 80 GB). **b,** The top 10 HVGs found across the spatial domains defined by the embeddings of different methods (SLAT, Banksy, DECIPHER); notably, CXCL13_CXCR5 was not found as an HVG through these methods (adjusted p values 0.39, 0.46 and 0.43, respectively) (**Methods**). **c,** Spatial expression patterns of CXCR4, CXCL12, CXCR5, and CXCL13 in B cells. **d,** Violin plot showing the LR activity of CXCL12_CXCR4 and CXCL13_CXCR5 in B cells (**Methods**).

**Extended Data Fig. 5:**
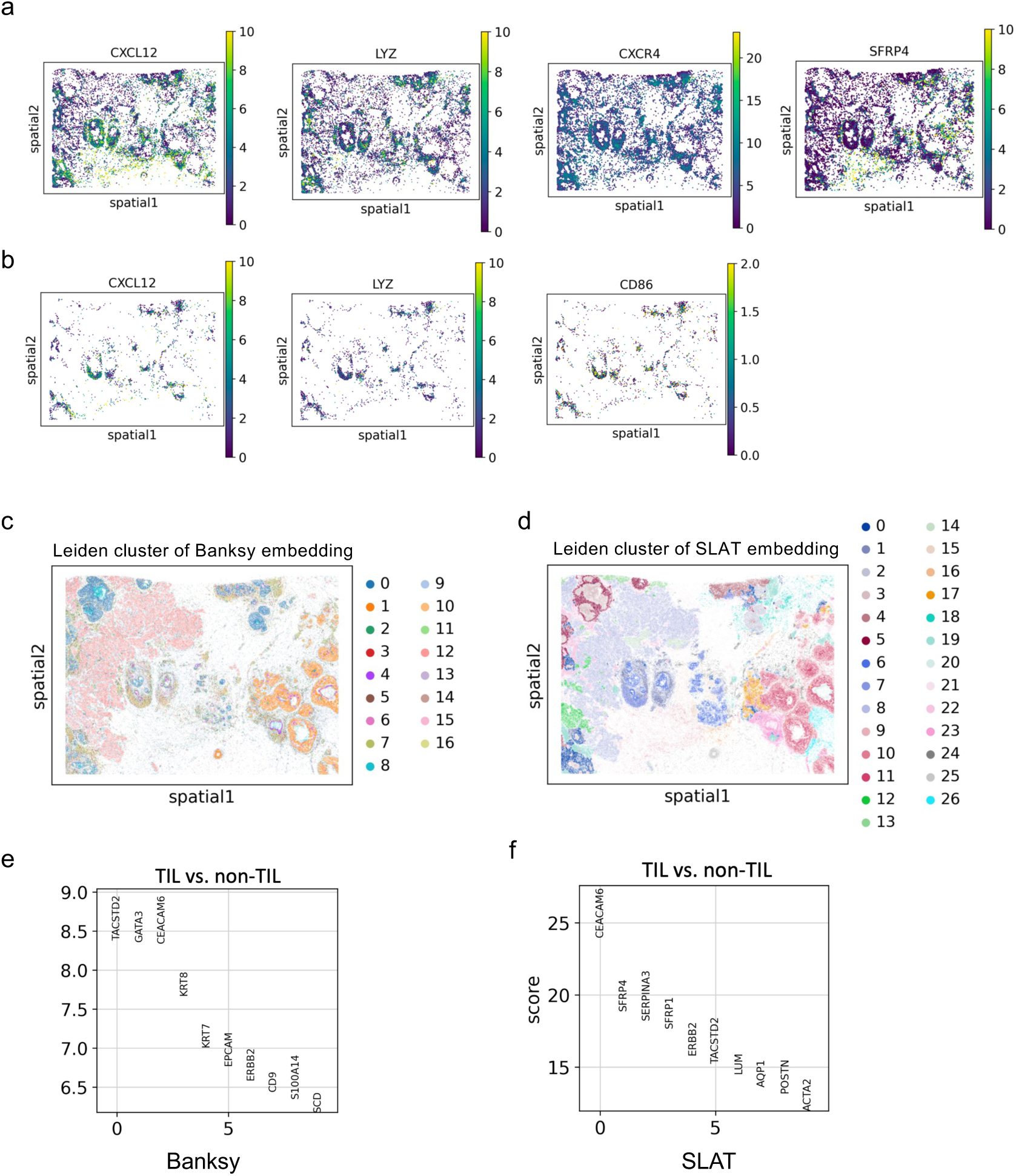
DECIPHER profiling of the breast cancer dataset. **a,** Spatial expression of the top 5 T cells and B cells (except PTGDS). **c, d**, Leiden clustering results based on Banksy embedding (**c**) and SLAT embedding (**d**). **e, f**, Top 10 HVGs identified between TILs and nontumor infiltrates lymphocytes (non-TILs) on the basis of the spatial domains defined by Banksy embedding (**e**) and SLAT embedding (**f**).

**Extended Data Fig. 6:**
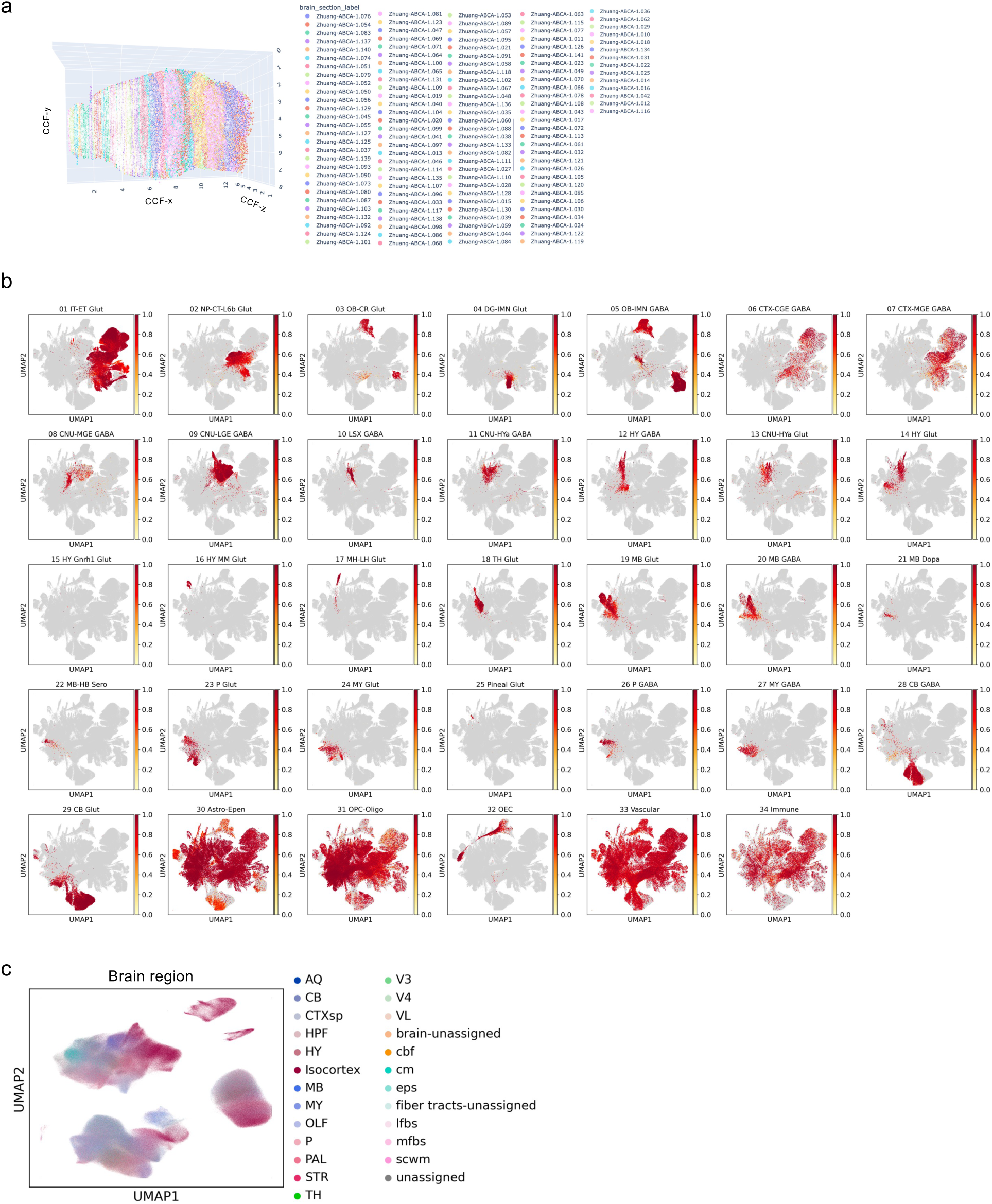
DECIPHER profiling of the 3D atlas of the mouse brain. **a**, Diagram illustrating each slice of the 3D mouse brain atlas. **b,** Distribution density of each cell type on the spatial context embedding UMAP (**Methods**). **d,** UMAP visualization of the DECIPHER spatial context embedding when only 2D cell coordinate information is used.

**Extended Data Fig. 7:**
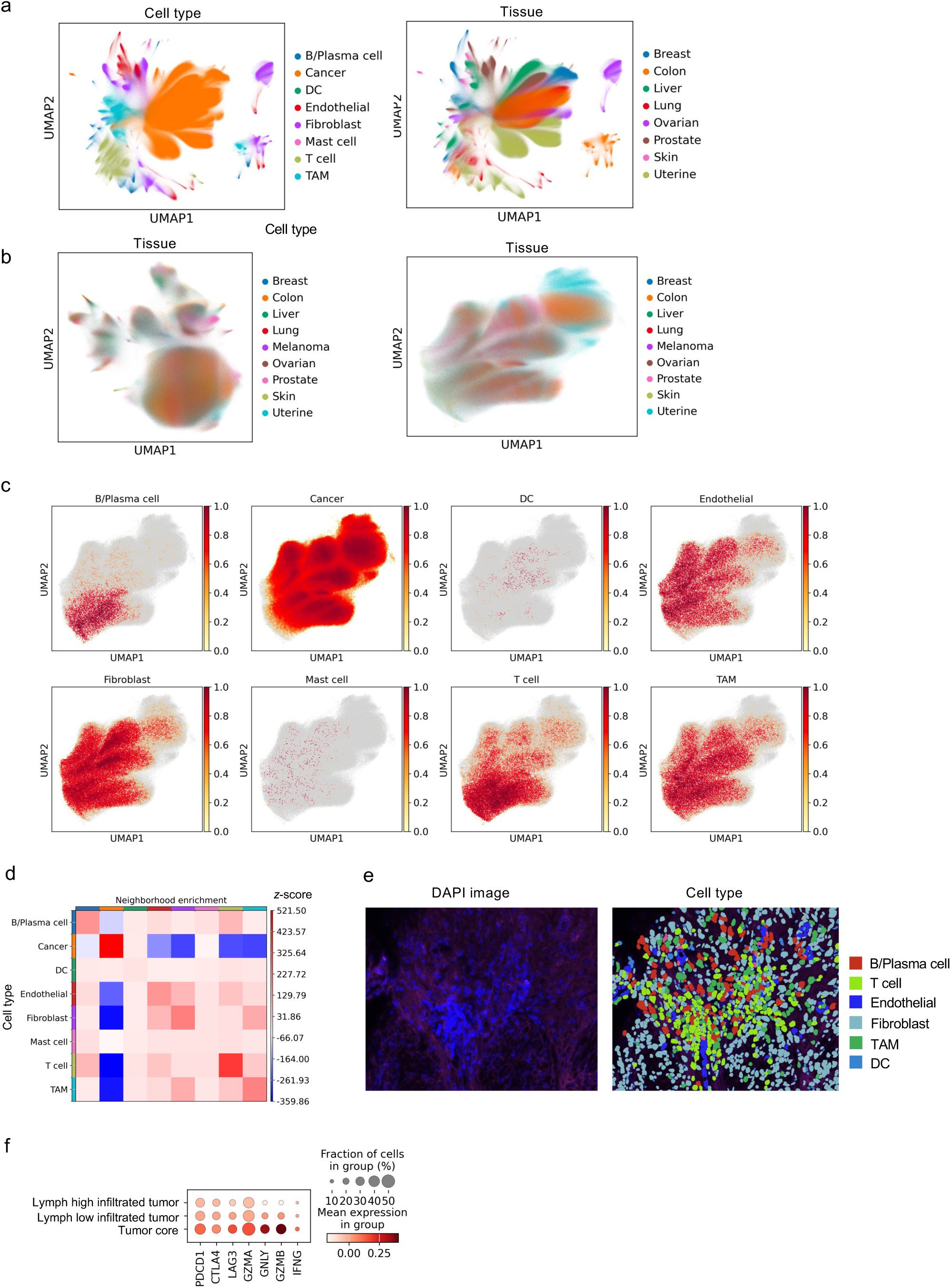
DECIPHER profiling of the human pan-cancer spatial atlas. **a, b,** UMAP visualization based on vanilla PCA embedding, colored by cell type (**a**) and tissue (**b)**, respectively. **c**, Distribution density of each cell type on spatial context embedding UMAP (**Methods**). **d**, Heatmap showing spatial enrichment at the cell type level; the value is the z score calculated via a permutation test^40^. **e,** Microscopy image of a typical TLS defined in the human pan-cancer dataset. The left panel shows the raw image, while the right panel shows the same view FOV colored by cell type based on cell segmentation. **f,** Dot plot showing the expression of exhaust marker genes^28^ (*PDCD1*, *CTLA4*, *LAG3*, *GZMA*, *GNLY*, *GZMB*, and *IFNG*) in T cells across different spatial regions.

**Extended Data Fig. 8:**
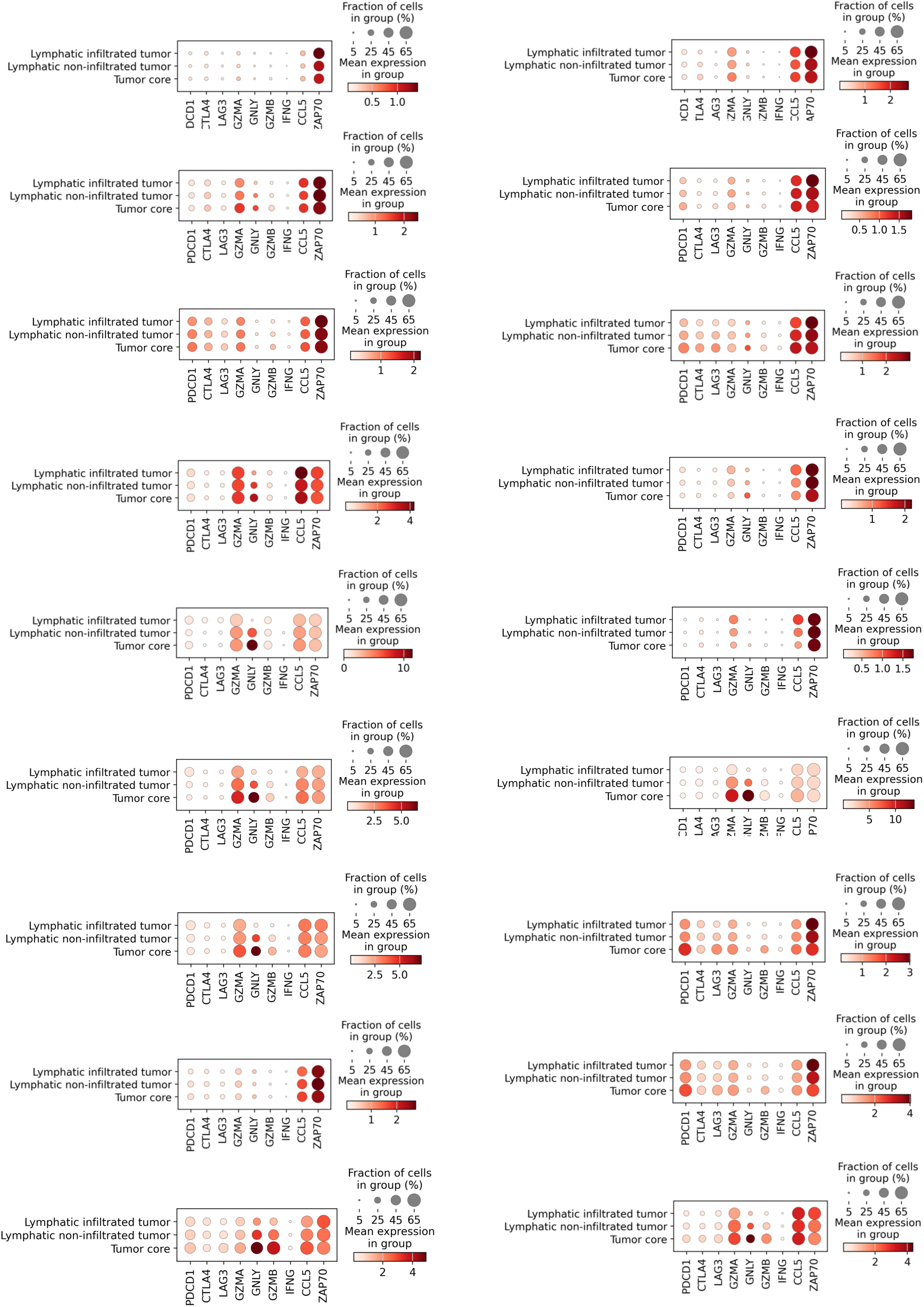
T-cell markers in each slice of the pan-cancer spatial atlas. Dot plot showing the expression of exhaust marker genes (*PDCD1*, *CTLA4*, *LAG3*, *GZMA*, *GNLY*, *GZMB*, and *IFNG*), *CCL5* and *ZAP70* in T cells across different spatial regions in each spatial slice of the human pan-cancer spatial atlas.

**Extended Data Fig. 9:**
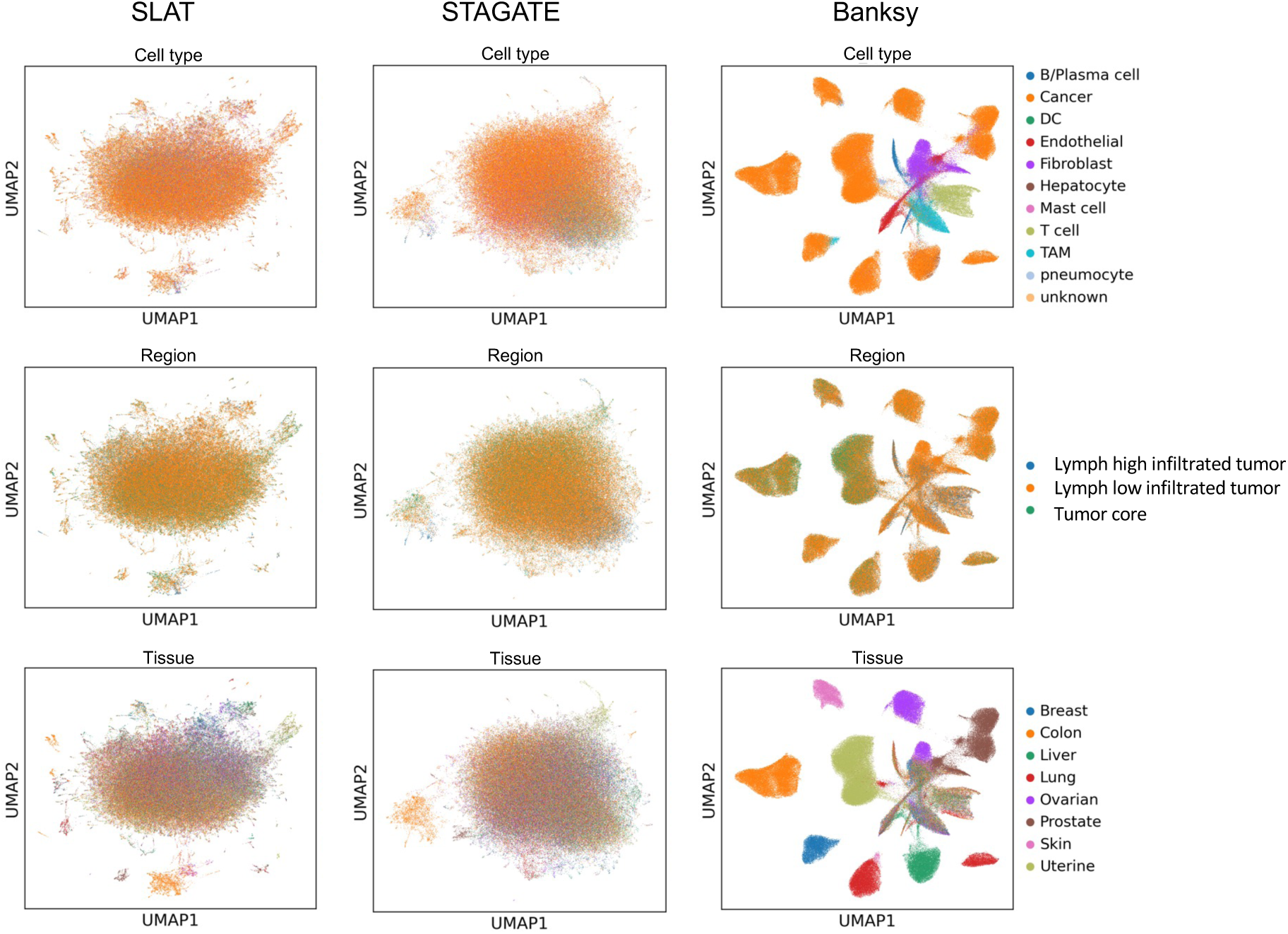
Comparison with other methods on down-sampled brain and pan-cancer atlases. UAMP visualization of SLAT, STAGATE and Banksy embeddings on a randomly down-sampled human pan-cancer spatial atlas with 200,000 cells, colored by cell type (first row), spatial region (second row) and tissue source (third row), respectively.

**Extended Data Fig. 10:**
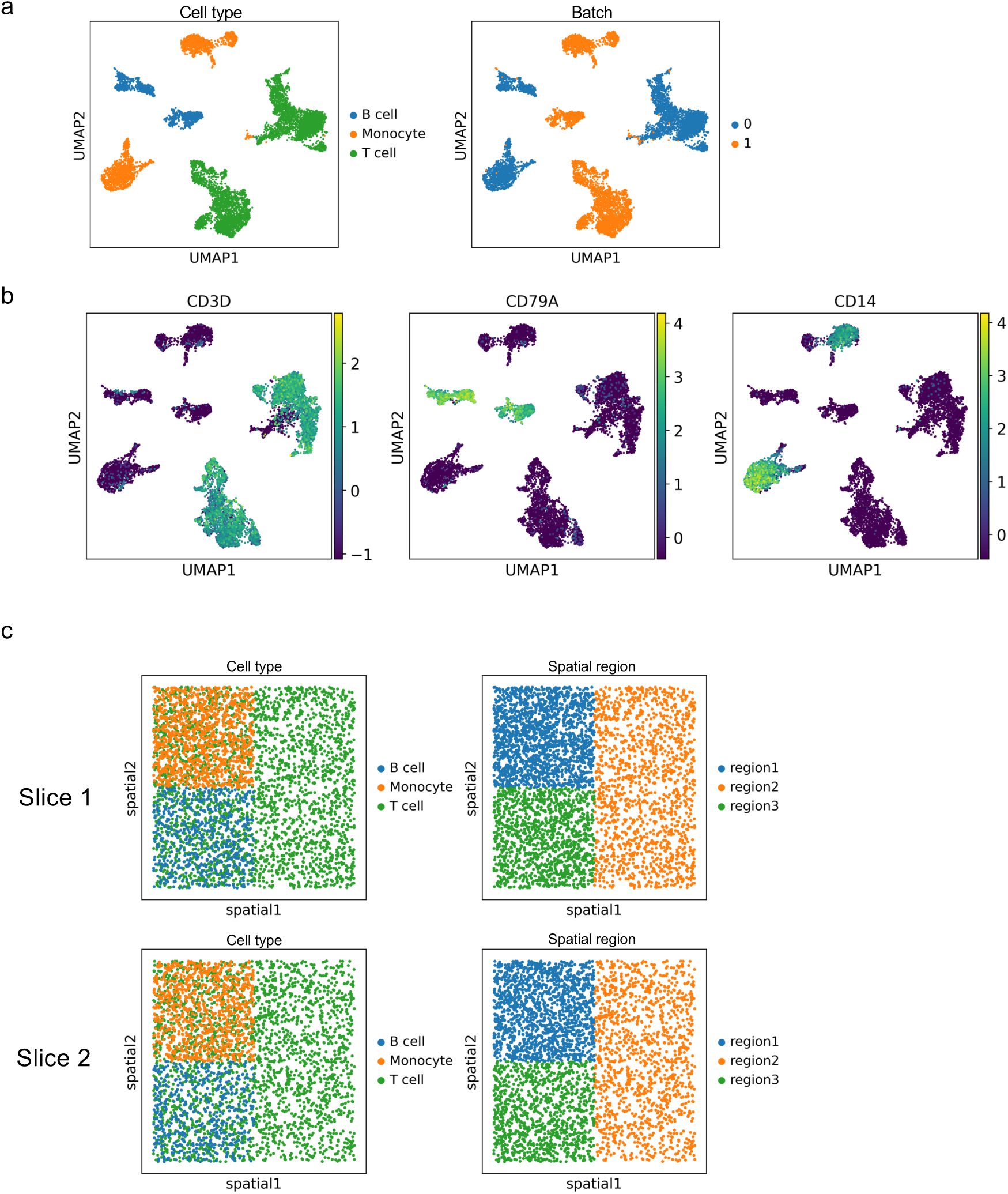
Simulation slices. **a,** UMAP visualization of PCA embeddings of two simulated slices, colored by cell type and batch. **b,** UMAP visualization of marker gene expression (*CD3D* for T cells, *CD14* for monocytes, and *CD79A* for B cells). **c,** Spatial visualization of simulated slice 1 (first row) and simulated slice 2 (second row), colored by cell type and spatial region, respectively.

